# Multiple imputation step-selection analysis: Improving estimation accuracy of travel distance accounting for route uncertainty

**DOI:** 10.64898/2026.02.23.707585

**Authors:** Shiori Takeshige, Jamie M Kass, Gota Yajima, Yusaku Ohkubo

## Abstract

1. Understanding animal movement behavior is essential for conservation and elucidating various ecological processes. In particular, assessing habitat suitability is a central theme in movement ecology, traditionally evaluated by estimating travel distances per unit time across diverse environmental conditions based on tracking data. Integrated step selection analysis (iSSA : Avgar et al., 2016) has been most widely applied in conservation studies to estimate animal movements that provide ecosystem services due to its ease of implementation and interpretability.
2. But despite its popularity, iSSA faces a critical issue—it can lead to underestimation of cumulative travel distances because it measures step lengths as conventional straight-line displacements between locations. This is primarily due to the application of linear interpolation between consecutive observed points, which fails to account for unobserved occurrences and non-linear trajectories taken by the individual.
3. In this paper, we propose a novel method to improve the estimation of travel distance in iSSA inspired by multiple imputation, a statistical method for missing data. We tested the extent to which our proposed method, Multiple Imputation Step Selection Analysis (MiSSA), improves the accuracy of step-length estimation (parameters of gamma distribution) compared to conventional iSSA using simulations across various scenarios. We compared the estimation bias and error under landscapes with different spatial autocorrelations (Simulation 1), as well as under various combinations of movement and habitat selection parameters (Simulation 2). In all simulations, MiSSA successfully reduced the estimation bias of the step length and substantially improved the estimation accuracy. In addition, although the coverage rates were comparable, we achieved a substantial reduction in the width of the confidence intervals.
4. Our study demonstrates that incorporating missing data statistics into the iSSA framework improves the accuracy of travel distance estimations, which serve as the foundation for evaluating habitat selection. MiSSA maintains the core advantages of iSSA while enabling more accurate estimation of travel distances, even for low-resolution data where movement between sampling intervals is non-linear. We anticipate its broad application across various disciplines, with a primary focus on conservation.

## 1. INTRODUCTION

Animal movement underlies many ecological processes, including population dynamics and species diversity across landscapes (Kays et al., 2015; Nathan et al., 2008). Understanding movement behavior enables us to identify key factors influencing dispersal, the core of metapopulation dynamics, as well as improve habitat connectivity and quantify the ecosystem services provided by animal movement (Baguette et al., 2013; Diniz et al., 2020; Earl & Zollner, 2017). Methodological development is essential to accurately measure and analyze movement behavior to better understand these ecological processes. In particular, biologging—the tracking of animals by attaching location-recording devices — has continued to advance and improve our understanding of animal movement behavior over the past decade (Nathan et al., 2022). These advancements have elucidated movement behavior that was previously difficult to record, such as travel routes, speed, and habitat selection (Watanabe & Papastamatiou, 2023), leading to rapid advances in understanding animal movement behavior. While most studies in this field have historically focused on large animals, an increasing number of studies now utilize tracking data of smaller mammals and birds (Nathan et al., 2022). However, as most devices available for tracking small animals (e.g., geolocators) have relatively longer positioning intervals, researchers must analyze movement patterns using data with relatively lower temporal resolution. When the sampling frequency is longer than the duration of the targeted behaviors (e.g., movement between migratory stopovers or within home-range movement), this hinders an accurate understanding of animal movement. Furthermore, it is often difficult to determine beforehand whether the sampling frequency aligns with the scale of the behavior being studied. Consequently, there is an urgent need to develop analytical methods that can draw inferences from these data across a wider range of taxa.

Integrated step selection analysis (iSSA) is one of the most frequently employed methods for movement data analysis (Avgar et al., 2016; Fieberg et al., 2021). iSSA is a statistical framework that simultaneously estimates habitat selection along with movement speed and path tortuosity (i.e., degree of meandering). It predicts the relative probability of selecting a step (movement from a location at time *t* to a location at time *t* + τ, where τ is the sampling frequency) by simultaneously estimating parameters for “selection-free” movement (i.e., a hypothetical context in which individuals show no preference for habitat) and those for habitat selection. These estimated parameters allow us to infer or predict the distributions of step length (travel distance between recorded locations) and turning angle (degree of rotation from the previous step), which form the basis for evaluating habitat selection. Because animals tend to move slowly and more tortuously in preferred landscapes, while moving quickly and linearly through unsuitable landscapes (Etzenhouser et al., 1998; Gillis & Nams, 1998; Odendaal et al., 1989), estimating these distributions across different spatial categories (i.e., land use) enables us to capture the characteristics of preferred versus avoided environments. iSSA has been widely applied in conservation, including identifying critical corridors (Hofmann et al., 2021, 2023), quantifying avoidance behavior in landscapes with strong anthropogenic impacts (Jansson et al., 2024; van Bemmelen et al., 2024), evaluating ecosystem functions provided by movement (Russo et al. 2024), and studying the habitat selection of reintroduced individuals (Majaliwa et al., 2022). iSSA has made significant contributions to understanding movement-driven ecological processes and will continue to play a central role in analyzing animal movement behavior.

But despite its usefulness, iSSA has a major limitation: it fails to account for the uncertainty between recorded locations and thus can underestimate travel distance. Although iSSA is a discrete-time model designed to evaluate movement at the temporal scale defined by the sampling interval without requiring the exact continuous trajectory, the estimated step length is inherently based on the straight-line minimum distance between observed locations.

Consequently, when the study focus is not only to estimate animal habitat selection but also to estimate true movement capacity by accounting for route uncertainty between location points, this straight-line approximation can inevitably lead to an underestimation of the actual travel distance when the temporal resolution of movement data is not sufficiently high. Although some modern tracking devices record location points at a higher resolution (e.g., one second) that can lead to more accurate movement-path data, several concerns remain. First, if the positioning frequency is too high, the measurement error of the tracking device brings significant noise, leading to an overestimation of travel distance (Noonan et al., 2019; Schoombie et al., 2024). Second, such devices are typically heavy and require recapture of subjects to collect recorded data, making them available only for limited species. In birds for example, attaching devices weighing more than 3% of an individual’s body mass is restricted for ethical reasons because it leads to loss of flight efficiency and reduced breeding success (Phillips et al., 2003). Availability of these devices can be further limited by behavioral traits; while these high-resolution devices typically require recapturing of tracked birds, this poses a difficulty unless they have a specific nesting location. Addressing these shortfalls would significantly help broaden the applicability of iSSA.

In this study, we propose a novel method using multiple imputation to improve estimates of travel distance with iSSA. Multiple imputation is a statistical method that predicts missing values in observed data by inserting plausible values to generate multiple “pseudo-complete” datasets that can be analyzed as if no data were missing (Rubin 1987). Several pioneering works in ecology have applied this technique in recent years, including Ohkubo et al. (2025) for regression analysis and Scharf et al. (2017) for movement data analysis. One recent study (Hoffman et al. 2024) found poor performance for an imputation approach for missing observations in movement data, but the single-state movement model they used for route interpolation might have limited capability in accurately reconstructing missing animal locations. Incorporating insights from the field of missing data analysis could improve the performance of travel distance estimation. Here, we introduce a method that imputes multiple values of step length and turn angles for estimating iSSA parameters, thus accounting for the uncertainty inherent in tracking data with low temporal resolution. To model animal movements from position *X_t_* to *X_t_*_+1_, we interpolate one position within this interval and calculate step length and turning angle. Then we obtain the maximum likelihood of coefficients for calculating the relative-use probabilities of each used and available step. This process is repeated multiple times (e.g., 100 to 1000) and different estimates are merged following Rubin (1987). While movement path has scale-dependency highlighted by fractal theory (e.g. Turchin 1996), our proposed method introduces a multiple imputation framework aimed at mitigating travel distance bias within the iSSA framework. In this paper, we first outline our proposed method for multiple imputation step-selection analysis, then describe two studies, one simulation and one empirical, to evaluate performance of this method. Finally, we discuss several limitations of this approach and potential future research.

## 2 PROPOSED METHODS

### 2.1 Overview of the multiple imputation step-selection analysis

In this section, we first provide an overview of the proposed model, which uses multiple imputation step-selection analysis (MiSSA) on tracking data to reflect uncertainty in routes between recorded location points (Fig. 1). Our main strategy to analyze GPS-recorded locations of animal movement is similar to previous methodologies, including iSSA. We use conditional logistic regression (1 for location use, 0 for otherwise) estimated with maximum likelihood to infer the relative probability that a location is used by an animal. MiSSA differs in that we introduce the novel concept of “imputed used steps”, which aims to calibrate the uncertainty of the route between observed locations. Imputed used steps are an extension to existing step-selection analyses that consider various possible routes through which animals can pass to mitigate travel distance bias in iSSA. In this method, all routes between location points are assumed to be missing data, and we substitute various routes to infer these missing routes based on multiple imputation. Finally, to estimate habitat selection and movement parameters, we integrate the imputation results based on the Rubin rule (Rubin, 1987), which we explain in 2.1.4.

**Figure 1.**
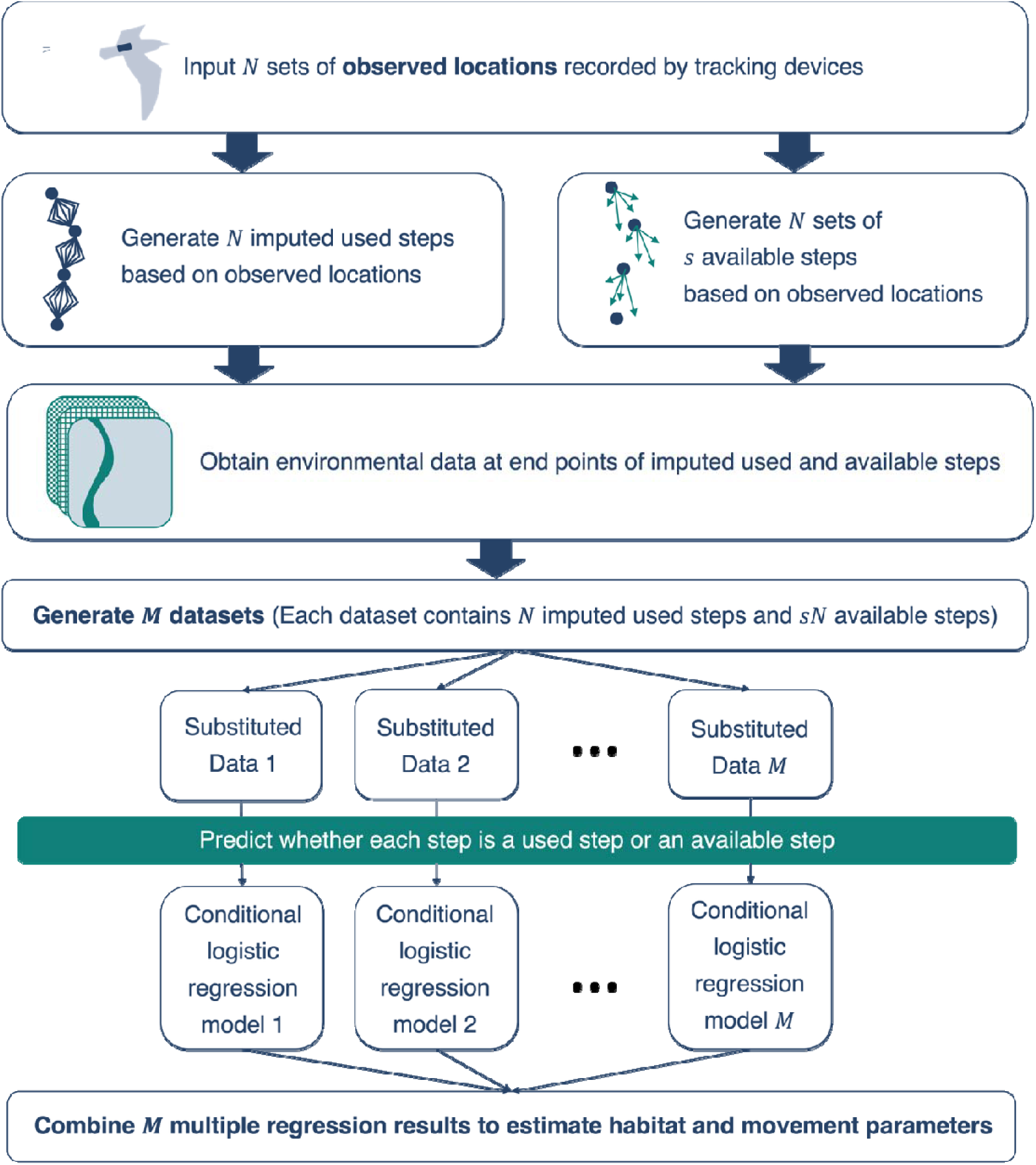
Flow chart depicting the workflow of multiple imputation step-selection analysis. First, data are preprocessed, then location data obtained by tracked animals are analyzed with MiSSA, where *M* is the number of multiple imputations, *N* is the number of imputed used steps contained in each dataset, and *s* is the number of available steps for each imputed used step. Unlike iSSA, this method involves imputing missing locations between observed locations *M* times and combining the results for parameter estimation. We give details of each procedure in the following sections.

#### 2.1.0 Notations and Previous Methods

Suppose we record the location *X_t_* in (*x_t_*, *y_t_*) of an animal at time *t* = 1, 2, … *T* with time interval τ. We assume each location is recorded without or with relatively low error, following previous studies using iSSA. Based on this data, we obtain three variables: *H*(*X_t_*_+*τ*_) is a set of environmental variables (e.g., elevation, land use, or distance from the nearest foraging site); *l_t_* is the step length (sl), which is the Euclidean distance from *X_t_*_−*τ*_ to *X_t_*; and *α_t_* is the turning angle (ta). The probability density of observing the location at *X_t_*, which is used by the animal at time *t* + τ, is modeled in conditional form as:

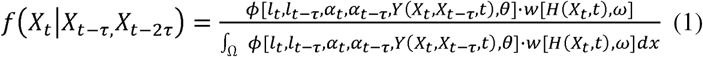

where the denominator is an integral over the entire spatial domain, Ω. iSSA is formulated by the “selection-free” movement kernel, *φ*, and the habitat selection function, *w* (eqn 1). This likelihood function poses practical difficulties since the denominator has an intractable integral. Thus, its approximation is often used to find a maximum likelihood estimate of the model, which reduces to the likelihood function of the conditional logistic model with environmental variables, step length, and cosine turning angles as explanatory variables.

#### 2.1.1 Imputed used step

The concept of imputed used steps extends the previous “used step” to reflect the uncertainty of missing locations within a movement dataset. These are locations that animals can travel in sampling interval τ and are generated by imputing missing locations between observed records. Suppose we have the recorded locations at time *t* and *t* + τ, and the recorded interval between them is not sufficiently short due to practical issues (e.g., GPS equipment with low battery capacity). In this example, we have data for *t* + 2τ due to our low temporal resolution, but we are interested in estimating the animal’s location at *t* + τ. Here, we propose to generate additional possible steps for every τ to incorporate the uncertainty in the route between observation locations with the following methodology (Fig. 2):

1. Draw a perpendicular bisector between the start point and an end point of a recorded step.
2. Generate candidate imputation points on the drawn bisector. For example, in our analysis, we generated a sequence of points at one-meter intervals on this line.
3. Recover movement paths, each of which connects the three points (i.e., start point of step, candidate imputed point, and endpoint). Then, step length and turn angle are calculated for each path. The paths connecting these three points are defined as the imputed used step.
4. If the step length of a recovered path exceeds a threshold (e.g., two standard deviations of the observed step length), optionally reject this candidate.
5. Select an imputed used step at random with probability proportional to the density of the step length and turn angle distributions, which generates available points.
6. Repeat 3-5 until *M* imputed used steps for every pair of successive observed locations are stored.

**Figure 2.**
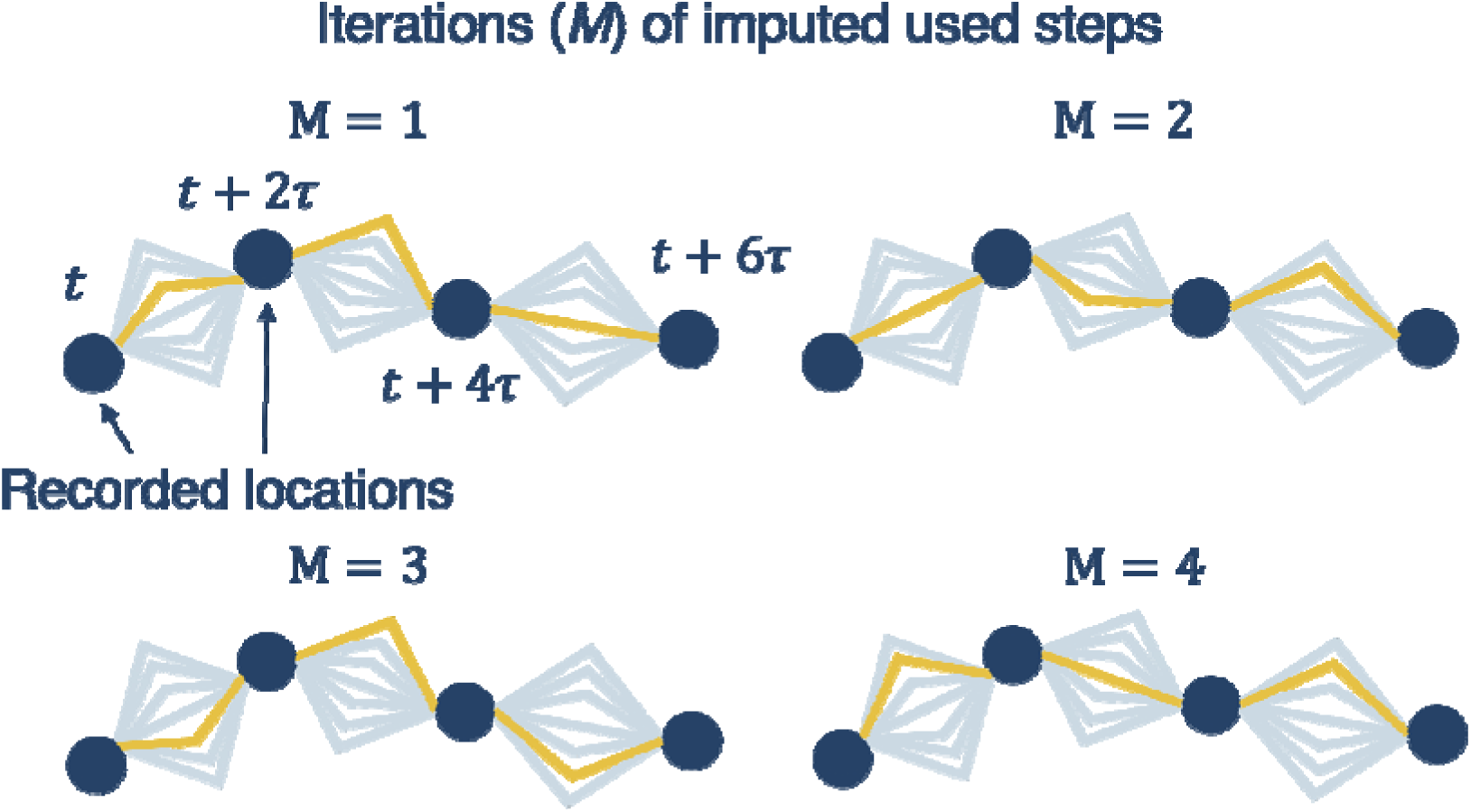
Illustrative example of imputed used steps. Suppose that the movement patterns of target animals can be better understood in τ = 1, although the observed data is recorded in τ = 2 owing to temporal resolution limitations of the tracking device. Imputed used steps represent the routes the animal is likely to travel during the sampling interval τ. Grey lines are candidate imputed used steps. Yellow lines are sampled imputed used steps. For every step *N* contained in the data, the imputed used step is sampled *M* times for multiple imputations. This figure shows the four different patterns of imputed datasets of (*M* = 4) for the same set of observed locations (*N* = 3).

We restricted the number of possible turns to one at the midpoint to form a perpendicular line due to computational constraints. Due to this restriction, we could not assume animal movements that turn multiple times or more than 90° away. However, animals tend to show highly directional movements while their migration. Therefore, the use of perpendicular lines in our method should describe such movements well. The fourth optional procedure is intended to remove less plausible routes, which reduces computational cost by limiting the number of possible patterns. Whether this step is needed and how the threshold is defined is open to the practitioner and would depend on the focus of the research question, as well as the quality and quantity of the data.

#### 2.1.2 Conditional Logistic Regression Analysis

Similar to the previous methods, we apply a conditional logistic regression model to predict the relative probability that a step is used and to estimate relationships with explanatory variables. The difference with previous methods is that we have *M* datasets (*i* = 1, 2, 3, …, *M*) with different imputed used steps, which are analyzed independently. The method for combining results from different datasets is explained in the subsequent section. For the *i*th dataset, the response variable for each step is a binary value: 1 for the imputed used steps, or steps for which the animal was actually present, and 0 for available steps, or steps assumed to be usable (similar to pseudoabsence data). The likelihood of the model is expressed as a function of regression parameters ***θ*** = (*θ_habitat_*, *θ_cos _ta_*, *θ_sl_*, *θ_log _sl_*)

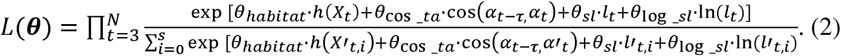

where ℎ(*X_t_*) is habitat value (continuous or discrete) at the end of step from *t* to *t* + *τ*, cos(*α_t_*_−*τ*,_*α_t_*) is the cosine of the direction of movement from *X_t_*_−*τ*_ to *X_t_* (turn angle), and *l_t_* is the Euclidean distance from *Xt*−*τ* to *Xt* (step length). Environmental conditions on the step can be reflected in habitat parameters *θ_habitat_* and step length and turn angles in movement parameters *θ_cos _ta_*, *θ_sl_*, *θ_log _sl_*. Each conditional logistic regression model uses *N* steps, the total number of *N* imputed used and *sN* available steps (*s* is the number of available steps per used step), i.e., *n* = *N* + *sN*, which is a vector of size *n* (*i* = 1, 2, …, *n*).

#### 2.1.3 Combining Multiple Results of Conditional Logistic Regression Analysis

Since MiSSA involves step selection analysis of multiple models, we need a formal rule to combine the different results of the regression analyses. We apply a rule inspired by the Rubin Rule, a method developed in the field of missing-data statistics (Rubin 1987). Our new route-selection model uses *M* conditional logistic regression models to predict whether each step is a used route or an available step, in contrast to existing step-selection analysis methods that use only one conditional logistic regression model. By analyzing each of these *M* datasets independently, we obtain different estimates of each coefficient θ̃ = (θ̃_1_, …, θ̃_*M*_). The within-imputation variance of θ̃ is shown in Equation 4 and its between-imputation variance is in Equation 5. To obtain the average effect of each parameter, the parameters obtained from the *M* conditional logistic regressions are merged following Rubin (1987) (Equations 3–6). Variance of θ̅_*M*_, *V_M_*, is calculated by integrating within-and between-imputation variance (Equation 6).

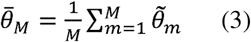

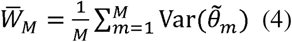

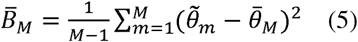

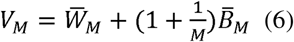

The relative probability of use of various steps under route uncertainty can be calculated using θ̅*_M_*, and confidence intervals can be calculated using *V_M_*. Movement parameters are updated using the combined coefficients calculated in eqn 3 as below.

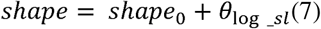

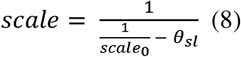

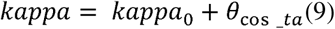

where *shape*_0_, *scale*_0_ and *kappa*_0_ are the parameters of step length and turn angle distributions, which generate available points.

## 3 Simulation Study

We conducted a simulation study to evaluate how the multiple imputed step selection analysis improves estimations of step lengths with iSSA. We generated animal movement paths in artificial landscapes and applied both the existing and proposed methods to predict step lengths. Then we compared the performance of the estimated step length with the true expected value. Simulations were performed in R version 4.4.2 (R Core Team 2025), though version 4.4.1 was also used to run tools in the NLMR package (details below) due to incompatibility issues.

### 3.1 Landscape simulation

We simulated virtual landscapes comprising continuous spatial layers with extents of 2000 × 2000 square pixels at 10 m resolution using a Gaussian random fields model implemented with the *nlm_gaussianfield* function in the package NLMR (Sciaini et al. 2018). We generated three landscapes with values for an arbitrary continuous environmental variable (e.g., elevation) with different maximum ranges of spatial autocorrelation (10, 15, and 20).

### 3.2 Movement simulation

To simulate movements across the virtual landscapes, we applied a simulation algorithm implemented in the package amt (Signer et al. 2019). The starting points of all simulated paths were set at random following a normal distribution with mean as center of the landscape and standard deviation as 300 m for both *x* and *y* coordinates. For landscapes with different maximum ranges of spatial autocorrelation (as described in 3.1), we set the true value of parameters with the *make_issf_model* function. In Simulation 1, we used the following settings: habitat parameter = 1; gamma distribution shape = 1 and scale = 50; and concentration parameter of von Mises distribution kappa = 0.5. In Simulation 2, for one landscape (maximum range of spatial autocorrelation = 10), we set various combinations of true parameters (Table 1). Then, the habitat value at the end of each step (*t* + *τ*) and movement kernels were simulated with the *redistribution_kernel* function and 1000 simulated paths each consisting of 20 steps at 2-minute intervals and 100 bursts (sets of steps) were generated.

**Table 1.** Six different sets of true parameters used to generate movement data for Simulation 2. We investigated the accuracy of the estimated parameters of the gamma distribution under six different sets of true parameters. For this simulation, spatial autocorrelation was set to 10. Habitat parameter is the value of an arbitrary continuous environmental variable (e.g., elevation). Shape and scale are parameters of the gamma distribution for step length. Kappa is a parameter of the von Mises distribution for turn angle. We generated 100 bursts (sets of steps), each consisting of 20 steps and ran a total of 1000 simulations.

| Set | Habitat<br>parameter | Shape | Scale | Kappa | Step number | Burst number |
| --- | --- | --- | --- | --- | --- | --- |
| Set 1 | 1 | 1 | 50 | 0.5 | 20 | 100 |
| Set 2 | 1 | 10 | 50 | 0.5 | 20 | 100 |
| Set 3 | 1 | 10 | 10 | 0.5 | 20 | 100 |
| Set 4 | 1 | 1 | 10 | 0.5 | 20 | 100 |
| Set 5 | 1 | 1 | 10 | 0.1 | 20 | 100 |
| Set 6 | 10 | 1 | 50 | 0.5 | 20 | 100 |

### 3.3 Fitting movement data to iSSA and MiSSA

We applied both iSSA and MiSSA to simulated movement data. While the simulated paths are generated with time interval τ, data are preprocessed to have time interval 2τ to emulate a low-resolution tracking device. Suppose a situation where the target animal’s behavioral characteristics are best captured at a sampling frequency of *τ*, yet tracking device constraints limit the maximum available resolution to 2*τ*. In such cases, it would be desirable to be able to capture the movement characteristics and habitat selection at the *τ* scale using only the available 2*τ* data. The number of available steps per (imputed) used step was set to 100 for both iSSA and MiSSA, with random steps generated by gamma and von Mises distributions. Initial parameters *shape*_0_, *scale*_0_ and *kappa*_0_ were estimated from the step lengths and the turning angles of the used steps without adjusting habitat environments, which are estimated using Euclidean distance and path tortuosity when recorded locations are linearly interpolated. We employed an approximation algorithm to calculate turning angles because the standard functions in the amt package required prohibitive computational time for our large-scale movement dataset. Upon performing ten validation trials for the parameter estimation, the total error across all parameters was found to be less than 10^-10^. In MiSSA, turn angle for each step was calculated as the average of the turn angles at the midpoint and endpoint of the step. Next, the habitat value (continuous variable) at the end point of each step was obtained. For iSSA, a conditional logistic regression model was used to estimate the coefficients for habitat, step length, log step length, and cosine turn angle. In contrast, MiSSA consists of three steps to estimate these coefficients. First, imputed used steps are generated between location points. Following 2.1.1-4, step length of imputed used steps was restricted. We weighted the imputed used steps using the same distribution as that employed in generating the available steps. Then, we sampled the imputed used steps between all location points for 100 multiple imputations. A relatively high number of imputations (e.g., 100 to 1000) is preferred to reduce simulation error (Royston & White 2011). Using 100 sets of imputed used steps and available steps, we estimated the coefficient values for each habitat variable, step length, log-transformed step length, and cosine turn angle in a conditional logistic regression model, yielding 100 coefficient values per each dataset. These different estimates for different datasets were then merged based on Rubin’s rule. Movement parameters (shape and scale in gamma distribution and kappa in von Mises distribution) are updated by using movement coefficients (step length, log step length and cosine turn angle) based on Equations 7–9.

### 3.4 Model evaluation

We compared the estimated step length distributions from both methods with the true values and evaluated how MiSSA improved the estimation of step length. To examine how landscape difference affects parameter estimation accuracy, we estimated gamma distribution parameters in three landscapes with different spatial autocorrelations (Simulation 1). In addition, we investigated the accuracy of the estimated parameter of the gamma distribution under six different sets of the true parameters (Simulation 2).

The true value of the step length is defined as the expected value of the gamma distribution, which is used to generate simulated movement paths. Note that simulated data are downsampled to have time interval = 2*τ*, rather than *τ*. According to the reproducibility of the gamma distribution, *sl_t_*_→*t*+*τ*_ + *sl_t_*_+*τ*→*t*+2*τ*_∼Gamma(*shape* + *shape*, *scale*) for any two Gamma distributed random variables *sl_t_*_→*t*+*τ*_∼Gamma(*shape*, *scale*) and *sl_t_*_+*τ*→*t*+2*τ*_∼Gamma(*shape*, *scale*) with a same scale parameter. We thus defined the true expected value as 2*shape* ∗ *scale* to evaluate accuracy of estimated distance assuming parameters are identical. In this context, reproducibility refers to the characteristic where, given two independent random variables sl_t→t+τ_ and sl_t+τ→t+2τ_, each distributed as Gamma(shape, scale), their sum sl_t→t+2τ_ is distributed as Gamma(shape + shape, scale).

Standard error of the expected value of the gamma distribution was estimated by Goodman’s (1960) variance of products formula:

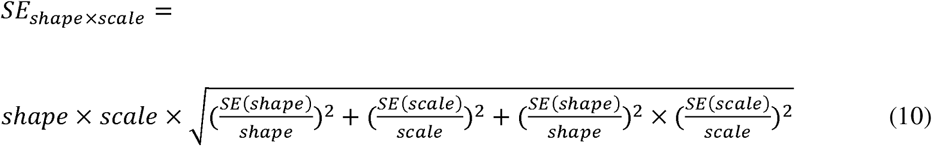

Based on the SE calculated using Equation 10, we compared the coverage probability of the expected value of the gamma distribution between iSSA and MiSSA. Because the sample size was sufficiently large, we calculated confidence intervals based on a normal approximation.

Furthermore, to assess the robust estimation of habitat selection under suboptimal sampling, we compared the true coefficients derived at frequency *τ* against the estimates obtained from iSSA and MiSSA using datasets thinned to 2*τ*. This comparison was designed to test if the models could recover “ideal” selection patterns from lower-resolution data. We conducted this comparison for both Simulation 1 and Simulation 2. Since the habitat parameter lacks the reproducibility that characterizes the gamma distribution, a direct comparison between the true values at frequency *τ* and the estimates from 2*τ*. is technically invalid. For this reason, we have presented these estimates as supplementary reference points.

## 4 Empirical case study

We applied MiSSA to empirical data and compared the results with iSSA. We used movement path data for the fisher (*Pekania pennanti*; individual Lupe) in the United States and landscape data (environmental variables) from Fieberg et al. (2021) downloaded via the amt package. A total of 14,229 GPS locations were recorded, with a median sampling interval of 2.1 minutes. The tentative Gamma and von Mises parameters were estimated as follows: shape = 1.44, scale = 36.27, and kappa = 0.49. We resampled the trajectory at 2-minute intervals (with a 20-second tolerance) using the *track_resample* function in R package amt, without applying any imputation methods for missing fixes. Following Fieberg et al. (2021), we assumed that habitat selection at the step scale depends on the three habitat variables. Land-use types were grouped into three categories: grass, forest, and wetland. Conditional logistic regression for iSSA and our model includes three habitat covariates, land-use types, elevation and population density at the end of each step and movement covariates (step length, log-transformed step length and cosine turn angle), as well as an indicator for the grouping that matches each used step to its corresponding available steps. As with the simulation study, in MiSSA, turn angle for each step was calculated as the average of the turn angles at the midpoint and endpoint of the step. We updated movement parameters based on Equations 7–9 and calculated the expected step lengths based on the shape and scale. Finally, we compared the mean squared error of the averaged value of the gamma distribution between iSSA and MiSSA.

## 1. Results

### 1.1 Simulation study

Under all simulation scenarios we tested, MiSSA yielded more accurate estimates of the expected gamma distribution value than iSSA (Table 2 and 3, Fig 4 and 5). Specifically, MiSSA was more accurate than iSSA for all landscapes with different levels of spatial autocorrelation in Simulation 1 and all parameter sets for Simulation 2.

**Table 2.** Comparisons of bias and coverage probability between iSSA and MiSSA in Simulation 1. The average of 1,000 step length estimates is reported for three different landscapes with maximum ranges of spatial autocorrelation spatial autocorrelation is 10, 15, or 20. We evaluated how closely the estimated mean expected value of the gamma distribution approximated the true expected value (2 × *shape* × *scale*). While iSSA underestimated the expected travel distance up to 15%, MiSSA overestimated the true expected value by only 3–4% in all landscapes we compared. The coverage probabilities across all models were 1.0, indicating that the 95% confidence intervals were too wide.

| Spatial autocorrelation | Model | True expected value of gamma distribution | Mean expected value of gamma distribution | Coverage probability |
| --- | --- | --- | --- | --- |
| 10 | iSSA | 100 | 84.15 | 1 |
|  | MiSSA |  | 103.98 | 1 |
| 15 | iSSA | 100 | 84.22 | 1 |
|  | MiSSA |  | 104.13 | 1 |
| 20 | iSSA | 100 | 84.21 | 1 |
|  | MiSSA |  | 104.05 | 1 |

In all scenarios we tested for Simulation 1, estimated values for MiSSA (103.98 - 104.13) were closer to the true value 100 than those for iSSA (84.15 - 84.22) (Table 2). In all landscapes, iSSA underestimated the expected value relative to MiSSA. This result showed that MiSSA reduced bias in estimation of the expected value of gamma distribution. In all landscapes, squared error was also smaller for MiSSA than iSSA (Fig. 3). Although coverage probability was overestimated in both methods (Table 2), estimation of confidence intervals improved with MiSSA (Fig. S1). Figure S2 presents the estimation results of the habitat parameter as a reference (note that the true values were based on a sampling frequency of *τ*, whereas the estimation used data sampled at 2*τ*), showing that this parameter was slightly overestimated by MiSSA. Neither method was able to accurately estimate kappa (Fig. S3).

**Figure 3.**
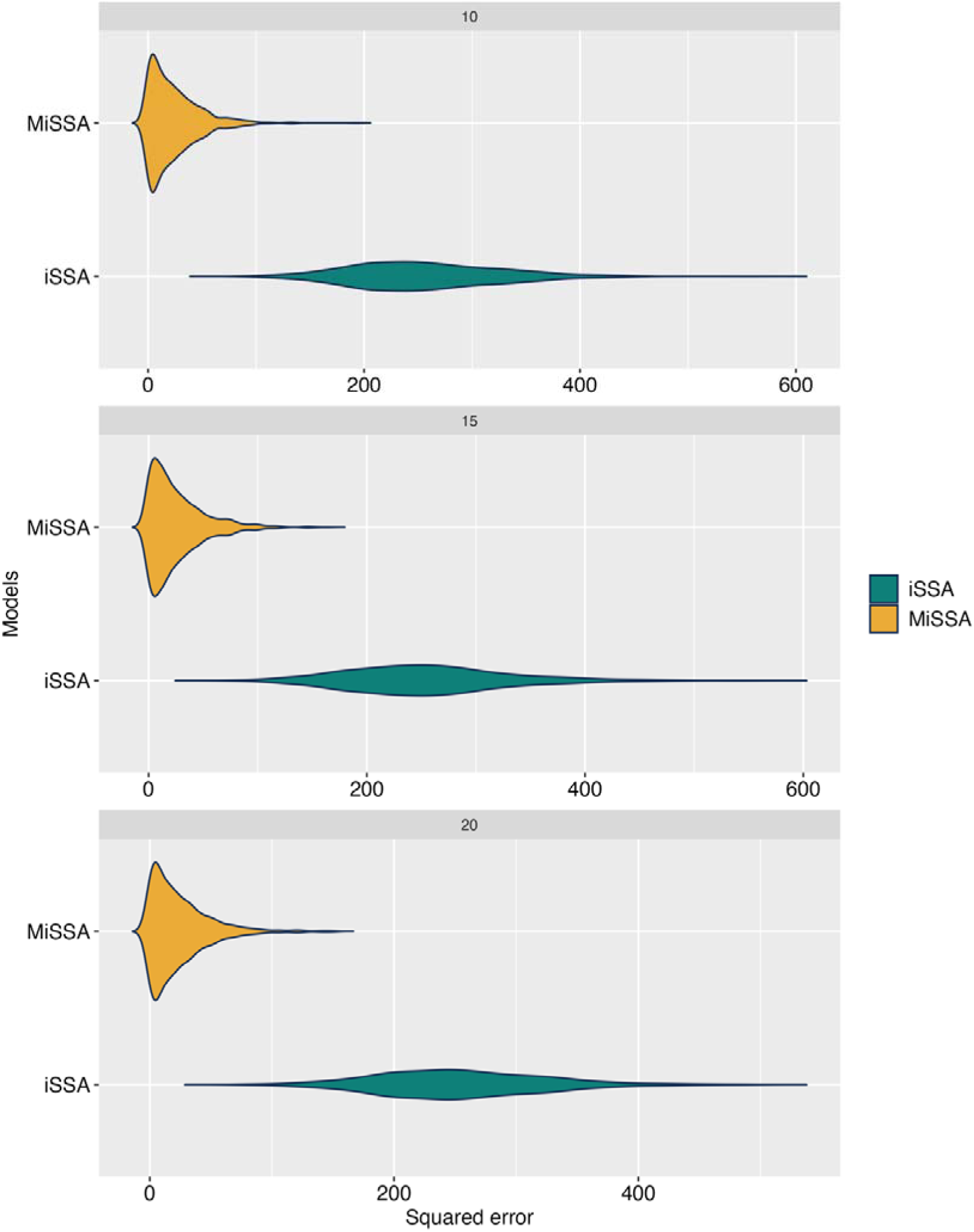
Comparisons of squared error between iSSA and MiSSA in Simulation 1. Violin plots depict squared error for estimations of the expected value of step length, obtained by 1000 replications of each simulation for different scenarios (three landscapes with different maximum ranges of spatial autocorrelation: 10, 15, and 20). In all landscapes, MiSSA had a smaller squared error and thus could estimate the expected value of the gamma distribution more accurately than iSSA.

For all parameter settings in Simulation 2, MiSSA was able to reduce the underestimation of the expected value of gamma distributions (Table 3). By accounting for path uncertainty between recorded locations, we mitigated estimation bias for the expected value of the gamma distribution within the iSSA framework (Fig. 4). For most parameter sets, the 95% confidence intervals for the expected value of the gamma distribution tended to be overestimated in both methods (Table 3). However, this tendency was much more pronounced in iSSA, indicating that MiSSA provided a higher estimation accuracy for these confidence intervals (Fig. S4). The estimation accuracy for the habitat parameters was almost identical for the two methods, although MiSSA slightly overestimated them in some parameter sets (Fig. S5). As for Simulation 1, in all parameter sets neither method was able to accurately estimate kappa (Fig. S6).

**Table 3.**
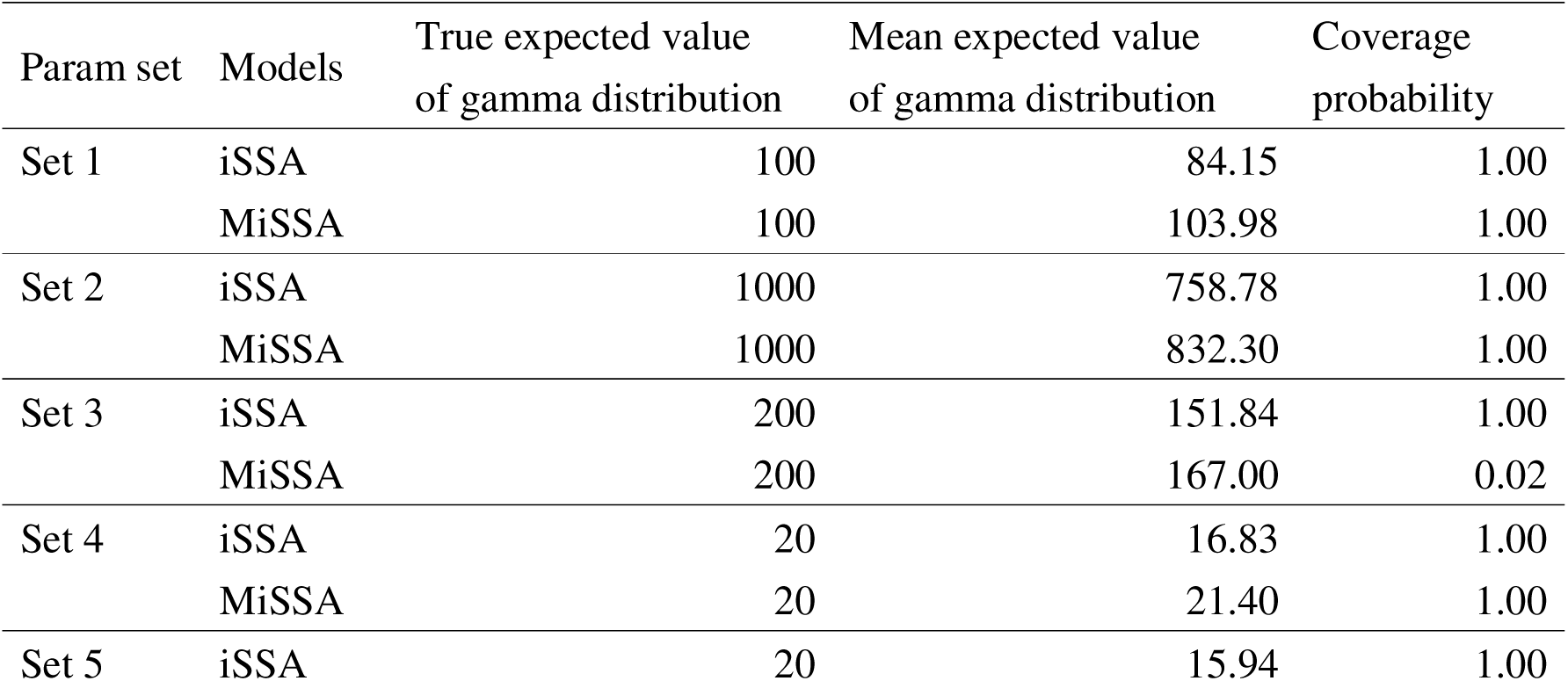
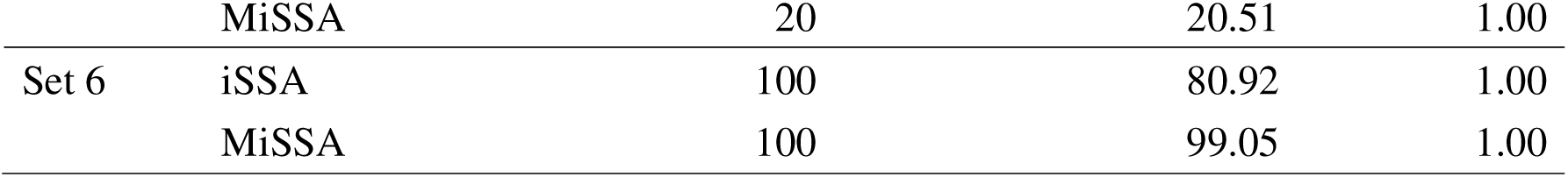
Comparisons of bias between iSSA and MiSSA in Simulation 2. We investigated the accuracy of the estimated expected value of the gamma distribution under six different sets of true parameters. Each of the six parameter sets comprises different gamma distribution parameters for step length, von Mises distribution parameters for turn angle, and habitat parameter. We then evaluated how closely the estimated mean expected value of the gamma distribution approximated the true expected value (2 × *shape* × *scale*). In all scenarios, MiSSA estimated the averaged value of the gamma distribution more accurately than iSSA, although the gap was less broad for a few scenarios (Sets 2 and 3).

| Param set | Models | True expected value<br>of gamma distribution | Mean expected value<br>of gamma distribution | Coverage<br>probability |
| --- | --- | --- | --- | --- |
| Set 1 | iSSA | 100 | 84.15 | 1.00 |
|  | MiSSA | 100 | 103.98 | 1.00 |
| Set 2 | iSSA | 1000 | 758.78 | 1.00 |
|  | MiSSA | 1000 | 832.30 | 1.00 |
| Set 3 | iSSA | 200 | 151.84 | 1.00 |
|  | MiSSA | 200 | 167.00 | 0.02 |
| Set 4 | iSSA | 20 | 16.83 | 1.00 |
|  | MiSSA | 20 | 21.40 | 1.00 |
| Set 5 | iSSA | 20 | 15.94 | 1.00 |
|  | MiSSA | 20 | 20.51 | 1.00 |
| Set 6 | iSSA | 100 | 80.92 | 1.00 |
|  | MiSSA | 100 | 99.05 | 1.00 |

**Figure 4.**
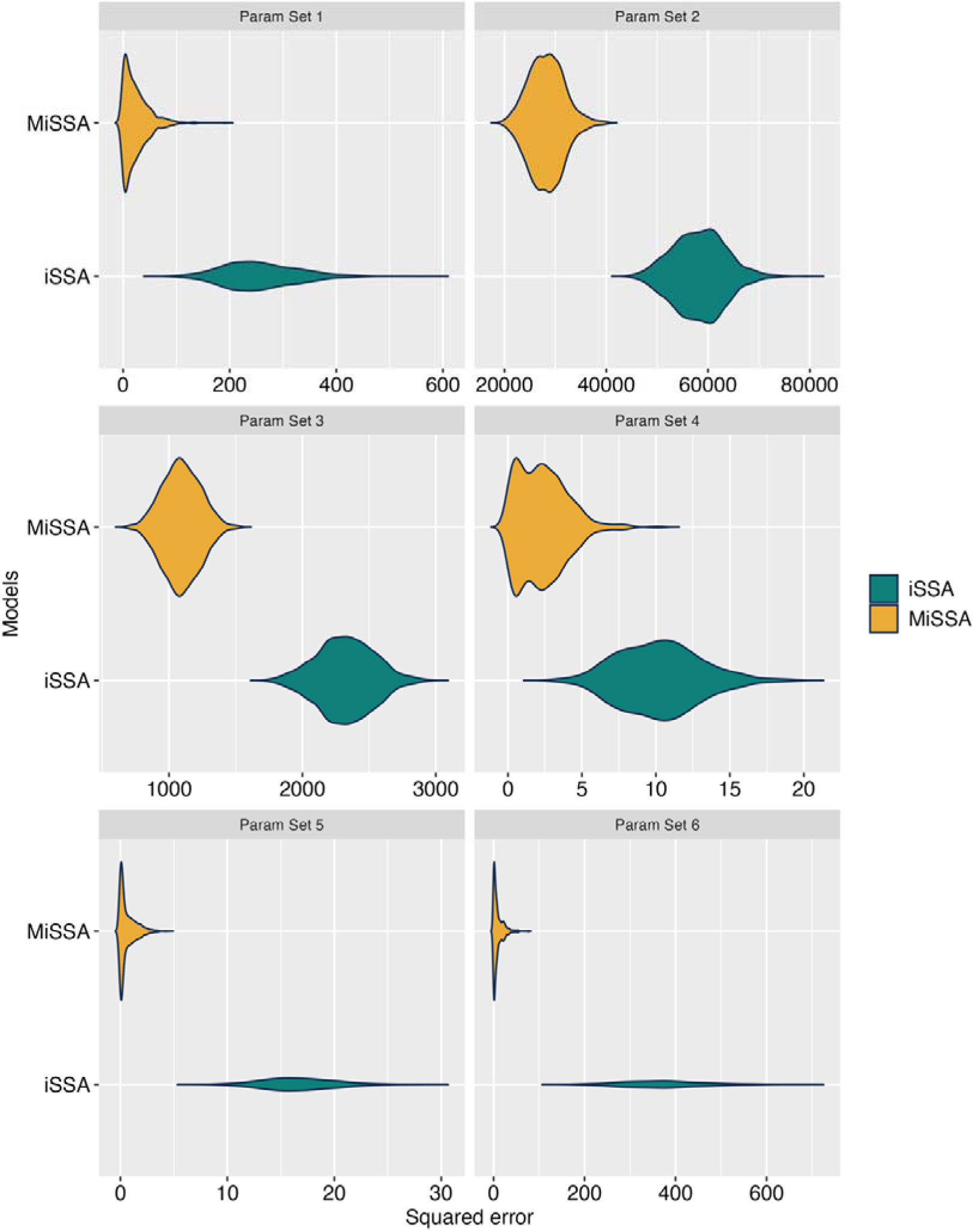
Comparisons of squared error between iSSA and MiSSA in Simulation 2.Violin plots depict squared error of step length estimates and are presented for six different parameter sets. To examine the accuracy of gamma distribution parameter estimation when the true parameters change, we estimated gamma distribution parameters using iSSA and MiSSA. In all scenarios, MiSSA had a smaller squared error and thus could estimate the expected value of the gamma distribution more accurately than iSSA.

Accuracy of estimated distance depended on the true value of parameter set, decreasing with higher continuous values of spatial, shape and scale parameters. In the scenarios with true parameters with high values (Sets 2 and 3), estimation bias and squared error increased (Table 3, Fig.4). Both methods underestimated the expected value of the gamma distribution in these scenarios; this underestimation was particularly notable in iSSA (Table 3).

### 5.2 Empirical case study

Consistent with the simulation results, the estimations of the expected gamma distribution values were lower for iSSA (53.48) than MiSSA (64.81) (Table 4). Thus, the relationship between the expected values for the two approaches aligned with the simulation results. As this is an empirical study. the true parameter values are unknown, yet we did observe that accounting for route uncertainty between recorded locations substantially influenced parameter estimation.

**Table 4.** Results of the case study for fisher (*Pekania pennanti*) tracking data. Estimated parameters are presented for habitat coefficients (population density, elevation and land use) and step length and turn angle distribution parameters were modeled by iSSA and MiSSA. Similar to the simulation results, the estimated expected value of the gamma distribution was larger for MiSSA than for iSSA.

| Model | Coefficients |  |  |  |  |  |  | Gamma distribution |  |
| --- | --- | --- | --- | --- | --- | --- | --- | --- | --- |
|  | Population density | Elevation | Land-use Grass | Land-use Wet | Shape | Scale | Kappa | Expected value (95% CI) |  |
| iSSA | -0.09 | 0.13 | -1.10 | 0.02 | 1.47 | 36.32 | 0.48 | 53.48 | (-57.14 - 164.10) |
| Proposed | -0.08 | 0.20 | -1.30 | 0.11 | 2.83 | 22.90 | 1.57 | 64.81 | (53.01 - 76.60) |

## 2. Discussion

In this paper, we propose a new method, multiple imputed step-selection analysis (MiSSA), to reduce bias in estimates of animal travel distance for movement ecology. Accurately modeling animal movement is essential for conservation and for understanding the ecosystem functions provided by animals, but traditional approaches to estimate step length do not account for uncertainty in routes caused by missing locations between observed location points. Most empirical studies have estimated average travel distance of tracking data by summing the straight-line distances between observed location points. While this method is straightforward and fast, it results in underestimation of the average travel distance when the sampling interval between location points has coarse temporal resolution, typically because animal movement between these points is not linear. In this study, we mitigated this issue by developing a novel imputation strategy for missing locations, and our results showed that it has higher performance than iSSA.

Specifically, by imputing multiple step lengths for various possible paths that animals can take between observed locations, we achieved more accurate estimation of the expected travel distance than iSSA. Although confidence intervals tended to be overestimated for both methods, MiSSA demonstrated an overall improvement in estimation accuracy. In cases where MiSSA randomly or excessively sampled detour locations, it would instead introduce a substantial positive bias, overestimating the true travel distance. While alternative configurations for constructing imputed used steps can certainly be envisioned—and identifying the absolute optimal approach remains a future challenge—our results demonstrate that even our parsimonious framework can profoundly improve the estimation accuracy of travel distances. The fact that MiSSA consistently reduces both estimation bias and squared errors proves that our corrections are well-bounded, stable, and statistically reliable. The fact that MiSSA consistently reduces both estimation bias and squared errors demonstrates that our corrections are well-bounded, stable, and statistically reliable. Related to this, although the temporal midpoint was selected to reduce computational costs, selecting alternative imputation points—such as the 1/3 or 2/3 marks—inherently would not change our core conclusion.

The use of imputation methods was questioned by Hofmann et al. (2024), who examined a variety of approaches to handle missing data within the iSSA framework and found that imputation methods performed poorly because of their single imputation approach. We avoided this problem by developing a multiple imputation approach, which should evaluate travel distance per unit time more accurately. Multiple imputation has been applied in previous research to incorporate uncertainties, such as location errors, into hidden Markov models that are commonly used for estimating behavioral states from tracking data (McClintock 2017), but this study is the first to integrate multiple imputation into iSSA. Another potential method is quantifying habitat selection and movement capacity from low-resolution data by using empirical correction factors derived from high-resolution data. Indeed, empirical correction factors derived from high-resolution trajectories are highly valuable for quantifying habitat selection and movement capacity from low-resolution data, but their practical application is inherently constrained by the feasibility of capturing fine-scale movement. For many taxa, particularly, migratory songbirds, obtaining the necessary high-resolution tracking data remains extremely difficult. MiSSA addresses this practical bottleneck by offering a robust analytical framework that mitigates travel distance underestimation based solely on available low-resolution datasets.

Accurate estimation of travel distance is essential for understanding animal movement behavior and for informing conservation priorities. Responses of travel distance per unit time to environmental factors indicate habitat selection preferences for individuals (Prokopenko et al., 2017; Scrafford et al., 2018; Thorsen et al., 2022). In landscapes modified by people that include features with high animal mortality like roads (Trombulak & Frissell, 2000), it is especially important to clarify average travel distances in order to ensure animal movement corridors are functional. For example, mortality risk can be high if travel distance per unit time increases as animals approach or cross a specific land-use (Dussault et al., 2007; Leblond et al., 2013). This information is essential to balance animal conservation and land-use management. Furthermore, by comparing average travel distances across different time windows and animal life stages, we can promote effective conservation measures tailored to specific seasons or times. Animal responses to human activities may vary depending on sex and life stage (e.g. (Ladle et al., 2019; Loggers et al., 2025). This can be clarified by calculating and comparing the average travel distance for each life stage and time window based on location data obtained from individuals. Therefore, accurate estimation of average travel distances can promote animal conservation measures, thereby reducing conflicts between animals and humans.

MiSSA also represents an advancement for making use of older tracking data, or those that due to various limitations can only be acquired at lower temporal resolutions. The utilization of these data poses a serious challenge to movement ecology for several reasons. First, older tracking data may be crucial for understanding the impact of disturbance on animal movement before and after the event, as coarse temporal resolution cannot be compared directly with new data collected at higher resolutions, and scaling one to match the other necessitates a loss of information. Since MiSSA is particularly useful for estimating travel distance where movement patterns (direction and speed) change on short timescales relative to the sampling interval, it enables us to compare habitat and movement parameters between data of different temporal resolutions, including those acquired in the future with sampling frequencies exceeding present capacity. Although MiSSA exhibited a slight bias in estimating habitat parameters, the direction of the habitat effects remained unaffected (i.e., reversing positive and negative effects), meaning this is likely a negligible concern for practical applications. Second, MiSSA can be useful in cases where the quality of tracking data is outside the control of the researchers. For example, meta-analyses that rely on previously registered data from databases such as Movebank (Kays et al. 2022) can lead to variation in recorded date and quality among samples. Utilizing existing data with low temporal resolution is essential to integrate various data sources in movement ecology.

Despite these advantages, however, MiSSA has several limitations to be solved in future studies. First, our methodology does not validate the accuracy of habitat selection coefficients, as our method aims to improve the estimation of travel distance. When estimating habitat selection coefficients and kappa (the von Mises distribution parameter) for data with reduced sampling intervals, it is impossible to derive the true values of the coefficients. This is because simulated routes were generated from parameters with different temporal scales, in contrast to those of the gamma distribution where the reproducibility applies. However, our results suggest that MiSSA can estimate habitat parameters with reliable accuracy (Figs. S2 and S5), although the true values cannot be directly and precisely compared with the data from shortened sampling intervals. In most studies using selection functions, the main focus is determining preference or avoidance (i.e., whether the effect is positive or negative). In light of this purpose, the reference accuracy of the habitat parameters obtained in this study should be sufficient for practical applications. As with the habitat parameters, kappa cannot be directly and accurately compared in terms of accuracy due to a lack of reproducibility. Nevertheless, judging from our results (Figs. S3 and S6), it is challenging for both methods to capture movements that are finer in scale than those reflected in the recorded data frequency. This finding highlights that researchers whose primary focus is the estimation of turning angles need to be more prudent in choosing their sampling intervals. Second, when the true value of the shape parameter is large, the averaged value of the step length was underestimated (Parameter set 2 and 3 in Table 3). This likely arises because the range of candidate imputed steps was narrower, making imputed step lengths shorter. This observation may indicate that the range and angular thresholds to generate candidate imputed used steps should be flexibly adjusted depending on the research subject (e.g., foraging trip or migration), although establishing methods for setting this range and account for the degree of tortuosity remains a future challenge. Lastly, in our simulation experiments, coverage probability deteriorated for one of our parameter sets. This is likely due to the imperfect match between the generative and estimation models, but such effects on coverage probability should be investigated further.

## Acknowledgements

We thank Dr. Keisuke Yano, Dr. Kazuhiro Katoh, Dr. Kiyohiro Ikeda, Dr. Hiromichi Suzuki and Dr. Masato Yamamichi for their insightful comments and fruitful discussions.

## Author Contributions

Shiori Takeshige: conceptualization (lead), formal analysis (lead), methodology (lead), software (equal), writing – original draft (lead). Jamie M Kass: methodology (equal), validation (equal). Gota Yajima: methodology (equal), validation (equal). Yusaku Ohkubo: supervision (lead), software (equal).

## References

1. Avgar, T., Potts, J. R., Lewis, M. A., & Boyce, M. S. (2016). Integrated step selection analysis: bridging the gap between resource selection and animal movement. Methods in Ecology and Evolution, 7, 619–630. 10.1111/2041-210X.12528

2. Baguette, M., Blanchet, S., Legrand, D., Stevens, V. M., & Turlure, C. (2013). Individual dispersal, landscape connectivity and ecological networks. Biological Reviews, 88, 310–326. 10.1111/brv.12000

3. Diniz, M. F., Cushman, S. A., Machado, R. B., & De Marco Júnior, P. (2020). Landscape connectivity modeling from the perspective of animal dispersal. Landscape Ecology, 35, 41–58. 10.1007/s10980-019-00935-3

4. Dussault, C., Ouellet, J.-P., Laurian, C., Courtois, R., Poulin, M., & Breton, L. (2007). Moose movement rates along highways and crossing probability models. The Journal of Wildlife Management, 71, 2338–2345. 10.2193/2006-499

5. Earl, J. E., & Zollner, P. A. (2017). Advancing research on animal-transported subsidies by integrating animal movement and ecosystem modelling. The Journal of Animal Ecology, 86, 987–997. 10.1111/1365-2656.12711

6. Etzenhouser, M. J., Owens, M. K., Spalinger, D. E., & Murden, S. B. (1998). Foraging behavior of browsing ruminants in a heterogeneous landscape. Landscape Ecology, 13, 55 –64. 10.1023/A:1007947405749

7. Fieberg, J., Signer, J., Smith, B., & Avgar, T. (2021). A “How to” guide for interpreting parameters in habitat-selection analyses. The Journal of Animal Ecology, 90, 1027–1043. 10.1111/1365-2656.13441

8. Gillis, E. A., & Nams, V. O. (1998). How red-backed voles find habitat patches. Canadian Journal of Zoology, 76, 791–794. 10.1139/z98-017

9. Goodman, L. A. (1960). On the exact variance of products. Journal of the American Statistical Association. 55, 708–713. 10.1080/01621459.1960.10483369

10. Gould, L. A., Manning, A. D., McGinness, H. M., & Hansen, B. D. (2024). A review of electronic devices for tracking small and medium migratory shorebirds. Animal Biotelemetry, 12, 1–10. 10.1186/s40317-024-00368-z

11. Hofmann, D. D., Behr, D. M., McNutt, J. W., Ozgul, A., & Cozzi, G. (2021). Bound within boundaries: Do protected areas cover movement corridors of their most mobile, protected species? The Journal of Applied Ecology, 58: 1133–1144.

12. Hofmann, D. D., Cozzi, G., McNutt, J. W., Ozgul, A., & Behr, D. M. (2023). A three-step approach for assessing landscape connectivity via simulated dispersal: African wild dog case study. Landscape Ecology, 38, 981–998. 10.1111/1365-2664.13868

13. Jansson, I., Parsons, A. W., Singh, N. J., Faust, L., Kissui, B. M., Mjingo, E. E., Sandström, C., & Spong, G. (2024). Coexistence from a lion’s perspective: Movements and habitat selection by African lions (*Panthera leo*) across a multi-use landscape. PloS One, 19, e0311178. 10.1371/journal.pone.0311178

14. Kays, R., Crofoot, M. C., Jetz, W., & Wikelski, M. (2015). Terrestrial animal tracking as an eye on life and planet. Science, 348, aaa2478. 10.1126/science.aaa2478

15. Kays, R., Davidson, S. C., Berger, M., Bohrer, G., Fiedler, W., Flack, A., Hirt, J., Hahn, C., Gauggel, D., Russell, B., Kölzsch, A., Lohr, A., Partecke, J., Quetting, M., Safi, K., Scharf, A., Schneider, G., Lang, I., Schaeuffelhut, F., … Wikelski, M. (2022). The Movebank system for studying global animal movement and demography. Methods in Ecology and Evolution, 13, 419–431. 10.1111/2041-210X.13767

16. Ladle, A., Avgar, T., Wheatley, M., Stenhouse, G. B., Nielsen, S. E., & Boyce, M. S. (2019). Grizzly bear response to spatio-temporal variability in human recreational activity. The Journal of Applied Ecology, 56, 375–386.

17. Leblond, M., Dussault, C., & Ouellet, J.-P. (2013). Avoidance of roads by large herbivores and its relation to disturbance intensity: Avoidance of roads and disturbance intensity. Journal of Zoology, 289, 32–40. 10.1111/1365-2664.13277

18. Loggers, E. A., Litt, A. R., Haroldson, M. A., Gunther, K. A., & van Manen, F. T. (2025). Female and male grizzly bears differ in their responses to low-intensity recreation in a protected area. The Journal of Wildlife Management, 89, e70068. 10.1002/jwmg.70068

19. Majaliwa, M. M., Hughey, L. F., Stabach, J. A., Songer, M., Whyle, K., Alhashmi, A. E. A., Al Remeithi, M., Pusey, R., Chaibo, H. A., Ngari Walsoumon, A., Hassan Hatcha, M., Wacher, T., Ngaba, C., Newby, J., Leimgruber, P., & Mertes, K. (2022). Experience and social factors influence movement and habitat selection in scimitar-horned oryx (*Oryx dammah*) reintroduced into Chad. Movement Ecology, 10, 47. 10.1186/s40462-022-00348-z

20. McClintock, B. T. (2017). Incorporating telemetry error into hidden Markov models of animal movement using multiple imputation. Journal of Agricultural, Biological and Environmental Statistics, 22, 249–269. 10.1007/s13253-017-0285-6

21. Nathan, R., Getz, W. M., Revilla, E., Holyoak, M., Kadmon, R., Saltz, D., & Smouse, P. E. (2008). A movement ecology paradigm for unifying organismal movement research. Proceedings of the National Academy of Sciences, 105, 19052–19059. 10.1073/pnas.0800375105

22. Nathan, R., Monk, C. T., Arlinghaus, R., Adam, T., Alós, J., Assaf, M., Baktoft, H., Beardsworth, C. E., Bertram, M. G., Bijleveld, A. I., Brodin, T., Brooks, J. L., Campos-Candela, A., Cooke, S. J., Gjelland, K. Ø., Gupte, P. R., Harel, R., Hellström, G., Jeltsch, F., … Jarić, I. (2022). Big-data approaches lead to an increased understanding of the ecology of animal movement. Science, 375, eabg1780. 10.1126/science.abg1780

23. Noonan, M. J., Fleming, C. H., Akre, T. S., Drescher-Lehman, J., Gurarie, E., Harrison, A.-L., Kays, R., & Calabrese, J. M. (2019). Scale-insensitive estimation of speed and distance traveled from animal tracking data. Movement Ecology, 7, 35. 10.1186/s40462-019-0177-1

24. Odendaal, F. J., Turchin, P., & Stermitz, F. R. (1989). Influence of host-plant density and male harassment on the distribution of female Euphydryas anicia (*Nymphalidae*). Oecologia, 78, 283–288. 10.1007/BF00377167

25. Ohkubo, Y., Takeda, O., & Uchida, K. (2025). Causal and predictive data analysis for conservation: Simulation based comparisons and a case study for detecting impact of artificial feeding on Eurasian red squirrel (*Sciurus vulgaris*). Ecology and Evolution, 15, e72142. 10.1002/ece3.72142

26. Phillips, R. A., Xavier, J. C., & Croxall, J. P. (2003). Effects of Satellite Transmitters on Albatrosses and Petrels. The Auk, 120, 1082–1090. 10.1093/auk/120.4.1082

27. Prokopenko, C. M., Boyce, M. S., & Avgar, T. (2017). Characterizing wildlife behavioural responses to roads using integrated step selection analysis. The Journal of Applied Ecology, 54, 470–479. 10.1111/1365-2664.12768

28. R Development Core Team (2026) R: a language and environment for statistical computing. R Foundation for Statistical Computing, https://www.r-project.org/

29. Royston, P., & White, I. R. (2011). Multiple imputation by chained equations (MICE): implementation in Stata. Journal of statistical software, 45, 1–20. 10.18637/jss.v045.i04

30. Rubin, D. (1987). Multiple Imputation for Nonresponse in Surveys. Wiley. 10.1002/9780470316696

31. Russo, N. J., Nshom, D. L., Ferraz, A., Barbier, N., Wikelski, M., Noonan, M. J., Ordway, E. M., Saatchi, S., & Smith, T. B. (2024). Three-dimensional vegetation structure drives patterns of seed dispersal by African hornbills. The Journal of Animal Ecology, 93, 1935–1946. 10.1111/1365-2656.14202

32. Scharf, H., Hooten, M. B., & Johnson, D. S. (2017). Imputation approaches for animal movement modeling. Journal of Agricultural, Biological, and Environmental Statistics, 22, 335–352. 10.1007/s13253-017-0294-5

33. Schoombie, S., Wilson, R. P., Ropert-Coudert, Y., Dilley, B. J., & Ryan, P. G. (2024). The efficiency of detecting seabird behaviour from movement patterns: the effect of sampling frequency on inferring movement metrics in Procellariiformes. Movement Ecology, 12, 59. 10.1186/s40462-024-00499-1

34. Sciaini, M., Fritsch, M., Scherer, C., & Simpkins, C. E. (2018). NLMR and landscapetools: An integrated environment for simulating and modifying neutral landscape models in R. Methods in Ecology and Evolution, 9, 2240–2248. 10.1111/2041-210X.13076

35. Scrafford, M. A., Avgar, T., Heeres, R., & Boyce, M. S. (2018). Roads elicit negative movement and habitat-selection responses by wolverines (*Gulo gulo luscus*). Behavioral Ecology, 29, 534–542. 10.1093/beheco/arx182

36. Signer, J., Fieberg, J., & Avgar, T. (2019). Animal movement tools (amt): R package for managing tracking data and conducting habitat selection analyses. Ecology and Evolution, 9, 880–890. 10.1002/ece3.4823

37. Thorsen, N. H., Hansen, J. E., Støen, O.-G., Kindberg, J., Zedrosser, A., & Frank, S. C. (2022). Movement and habitat selection of a large carnivore in response to human infrastructure differs by life stage. Movement Ecology, 10, 52. 10.1186/s40462-022-00349-y

38. Trombulak, S. C., and Frissell, C. A. (2000). Review of ecological effects of roads on terrestrial and aquatic communities. Conservation Biology, 14, 18–30. 10.1046/j.1523-1739.2000.99084.x

39. Turchin, P. (1996). Fractal analyses of animal movement: a critique. Ecology, 77, 2086–2090. 10.2307/2265702

40. van Bemmelen, R. S. A., Leemans, J. J., Collier, M. P., Green, R. M. W., Middelveld, R. P., Thaxter, C. B., & Fijn, R. C. (2024). Avoidance of offshore wind farms by Sandwich Terns increases with turbine density. Ornithological Applications, 126, 5. 10.1093/ornithapp/duad055

41. Watanabe, Y. Y., & Papastamatiou, Y. P. (2023). Biologging and biotelemetry: Tools for understanding the lives and environments of marine animals. Annual Review of Animal Biosciences, 11, 247–267. 10.1146/annurev-animal-050322-073657

