## Supporting Information for "Multiple imputation step-selection analysis: Improving estimation accuracy of travel distance accounting for route uncertainty"

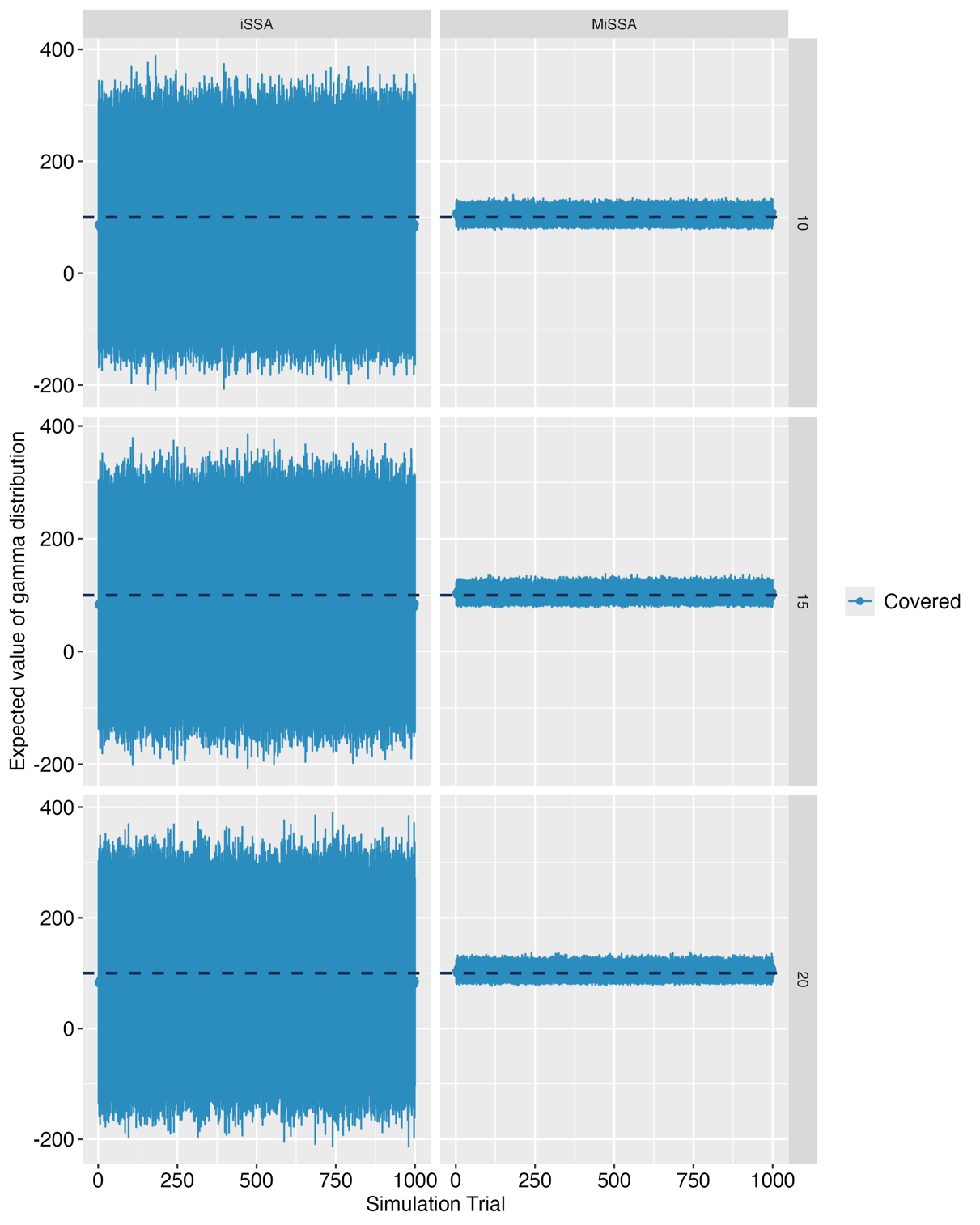

**Fig. S1** 95% confidence intervals for the expected value of the gamma distribution in simulation 1. Across all spatial autocorrelation landscapes (10–20), the 95% confidence intervals across all simulations covered the true expected value. However, the confidence intervals for iSSA were substantially wider than those for MiSSA.

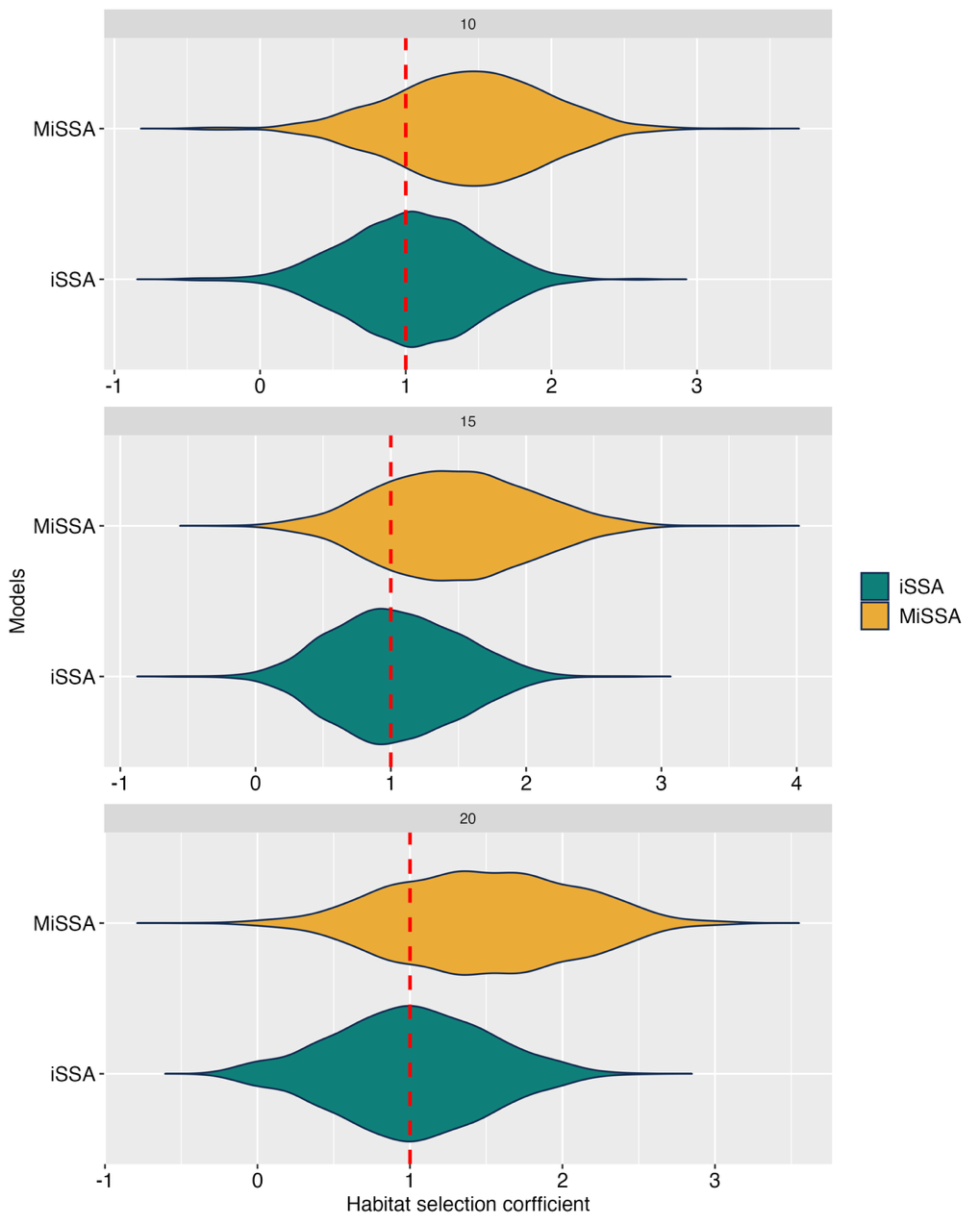

**Fig. S2** Comparisons of a habitat selection coefficient between iSSA and MiSSA in Simulation 1. Violin plots depict the value of habitat selection coefficient for different scenarios (three landscapes with different maximum ranges of spatial autocorrelation 10, 15, and 20). We evaluated how accurately the true value of $\Delta t$ could be estimated from the data thinned out to $\Delta2t$. The results indicated that while the MiSSA estimates exhibited a slight bias, they captured the same direction of effect.

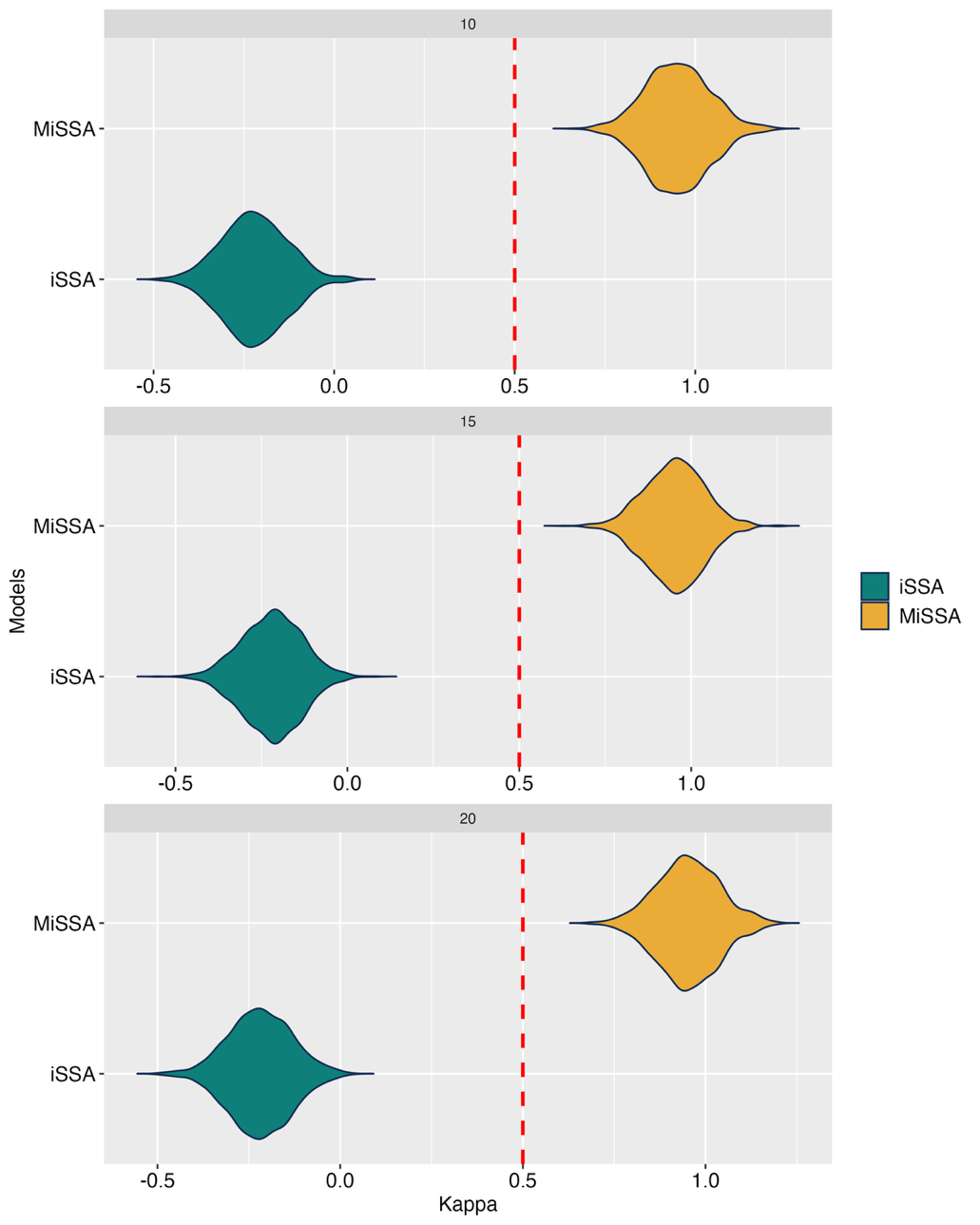

**Fig. S3** Comparisons of kappa parameter between iSSA and MiSSA in Simulation 1. Violin plots depict the value of kappa parameter for different scenarios (three landscapes with different maximum ranges of spatial autocorrelation 10, 15, and 20). We evaluated how accurately the true value of $\Delta t$ could be estimated from the data thinned out to $\Delta2t$. For both methods, the estimated values deviated from the true values because of the difference in sampling frequencies.

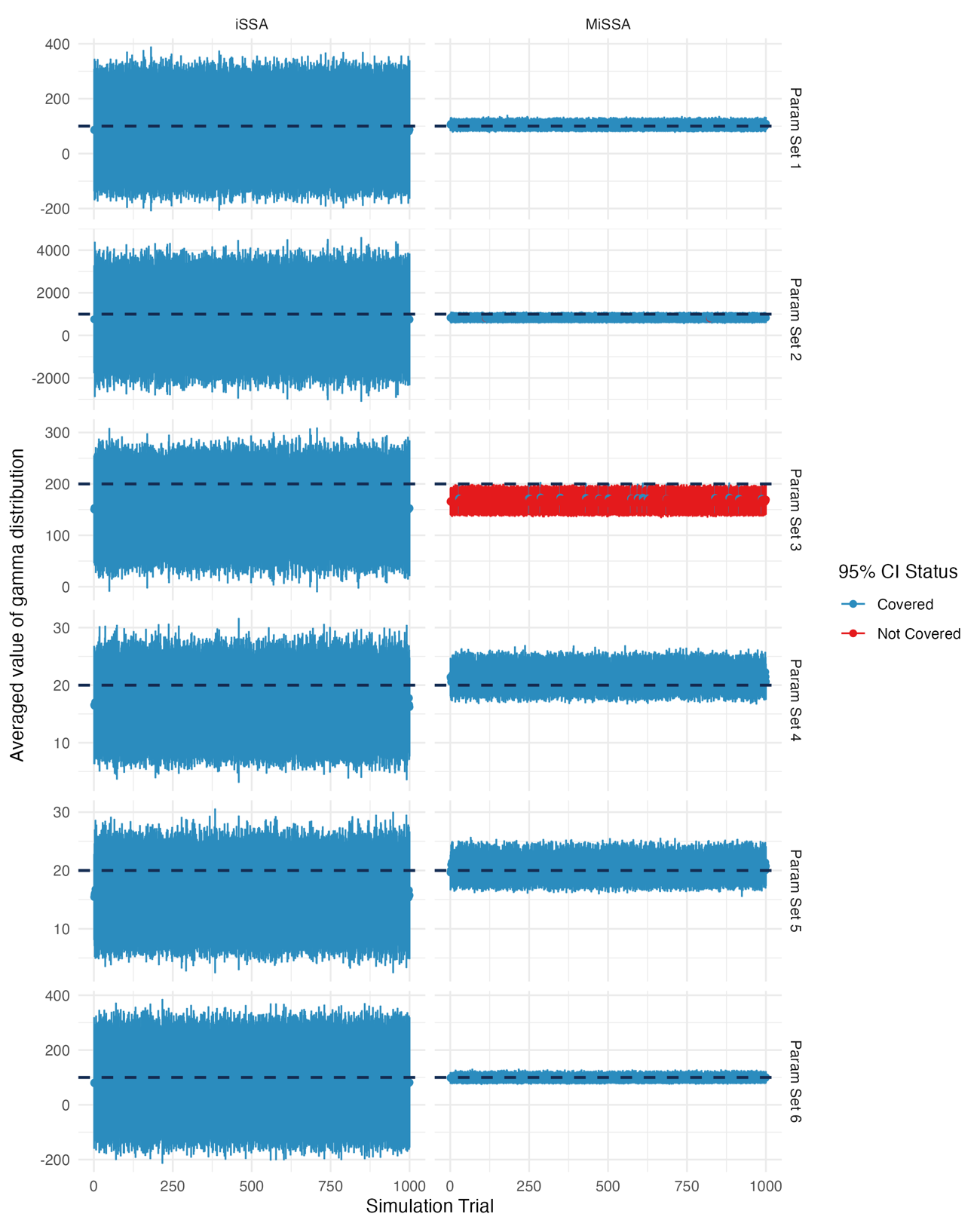

**Fig. S4** 95% confidence intervals for the expected value of the gamma distribution in simulation 2. In most parameter sets, the 95% confidence intervals across all simulations covered the true expected value. The confidence intervals for iSSA, however, were substantially wider than those for MiSSA. In parameter set 3, the coverage probability of MiSSA was considerably low, but its confidence interval width was much narrower than that of iSSA.

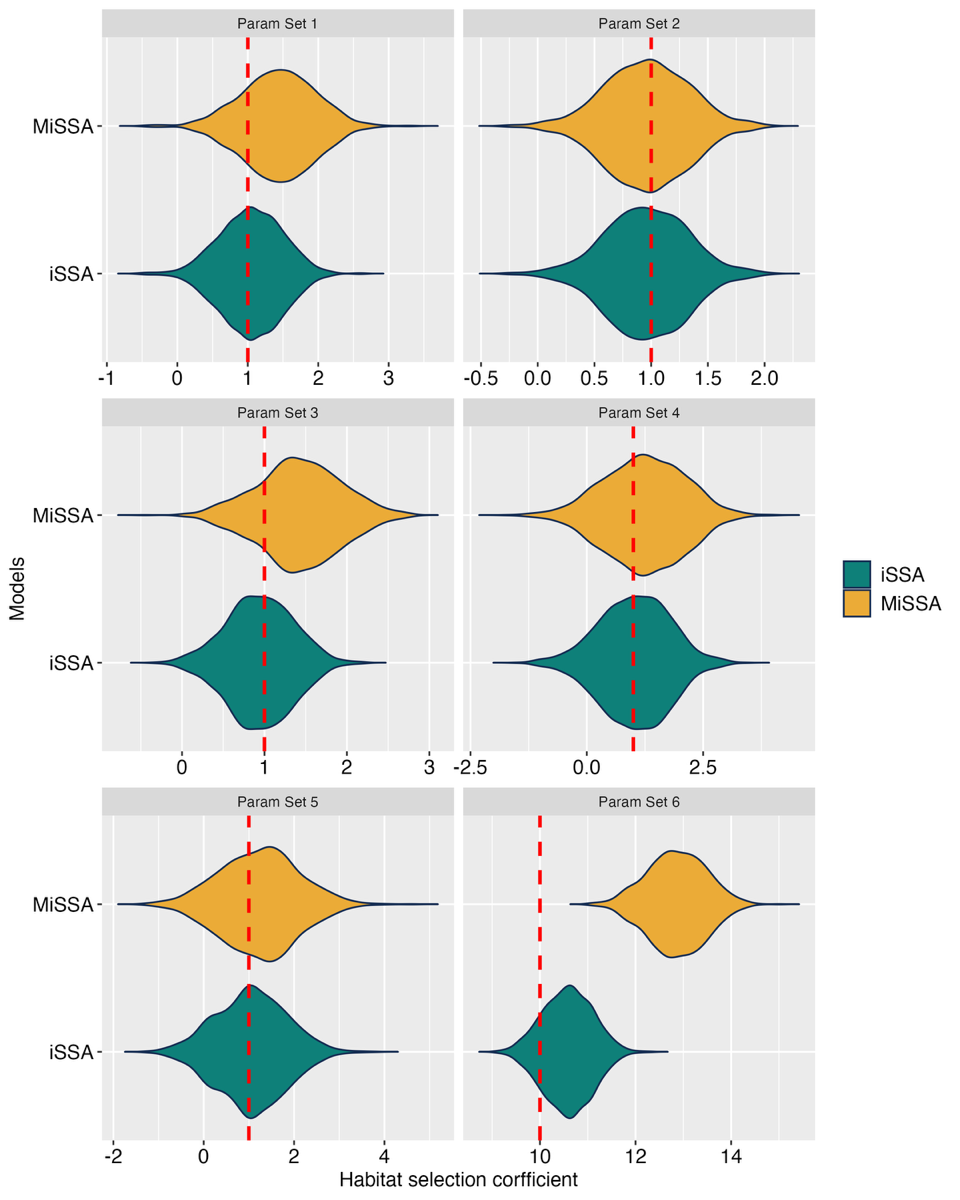

**Fig. S5** Comparisons of a habitat selection coefficient between iSSA and MiSSA in Simulation 2. Violin plots depict the value of habitat selection coefficient for different scenarios (parameter set 1-6). For parameter sets 1–5 (i.e., scenarios where the habitat selection coefficient equals 1), the results of iSSA and MiSSA were similar, and both methods provided comparably accurate estimates. However, for parameter set 6, which represents an extreme preference scenario with a habitat selection coefficient of 10, the estimates from both methods were biased.

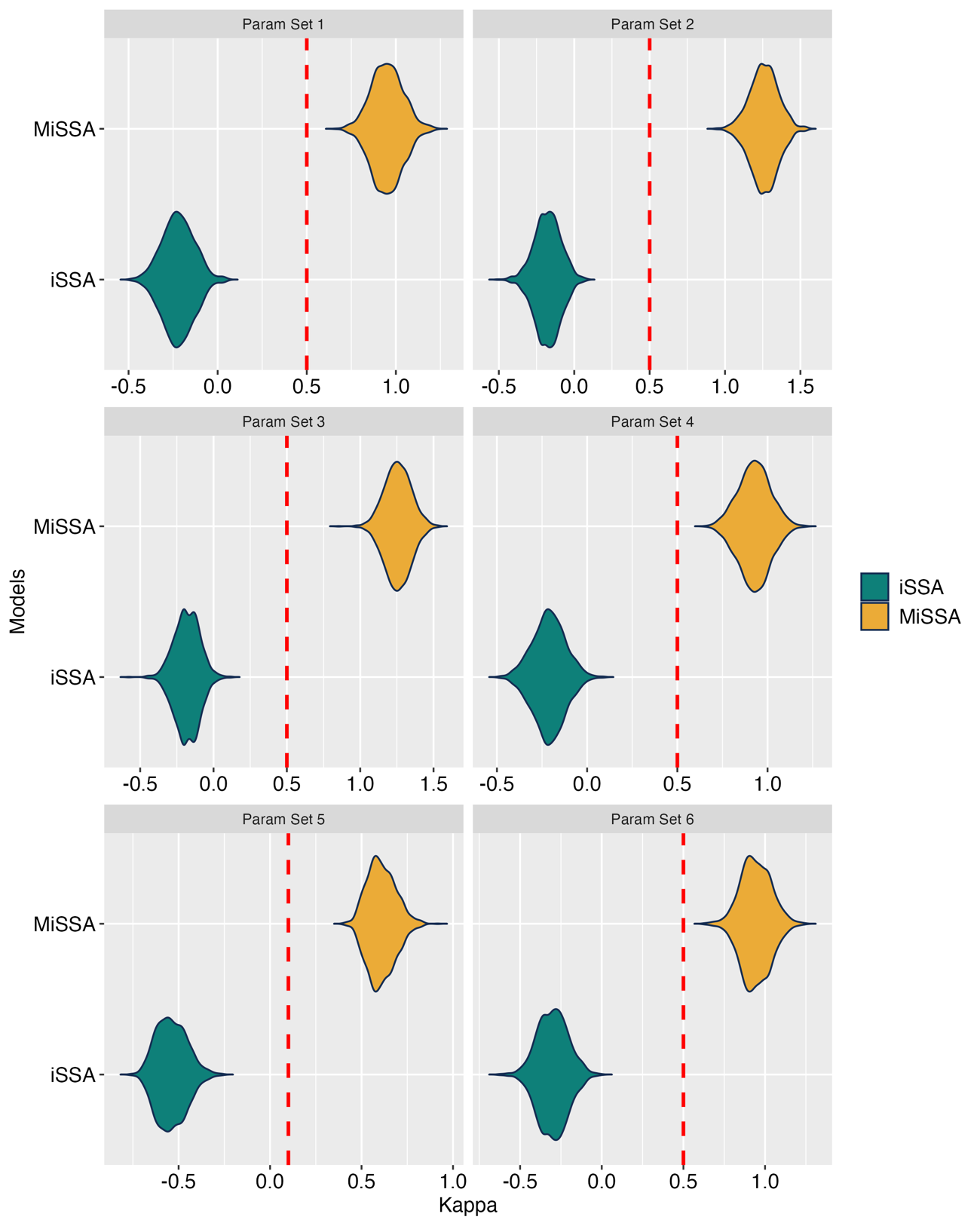

**Fig. S6** Comparisons of kappa parameter between iSSA and MiSSA in Simulation 2. Consistent with the results of Simulation 1, both methods failed to accurately estimate kappa across all parameter sets.
